# The Basic Reproductive Number for Disease Systems with Multiple Coupled Heterogeneities

**DOI:** 10.1101/220004

**Authors:** Alun L. Lloyd, Uriel Kitron, T. Alex Perkins, Gonzalo M. Vazquez-Prokopec, Lance A. Waller

## Abstract

In mathematical epidemiology, a well-known formula describes the impact of heterogeneity on the basic reproductive number, *R*_0_, for situations in which transmission is separable and for which there is one source of variation in susceptibility and one source of variation in infectiousness. This formula is written in terms of the magnitudes of the heterogeneities, as quantified by their coefficients of variation, and the correlation between them. A natural question to ask is whether analogous results apply when there are multiple sources of variation in susceptibility and/or infectiousness. In this paper we demonstrate that with three or more coupled heterogeneities, *R*_0_ under separable transmission depends on details of the distribution of the heterogeneities in a way that is not seen in the well-known simpler situation. We provide explicit formulae for the cases of multivariate normal and multivariate log-normal distributions, showing that *R*_0_ can again be expressed in terms of the magnitudes of the heterogeneities and the pairwise correlations between them. The formulae, however, differ between the two multivariate distributions, demonstrating that no formula of this type applies generally when there are three or more coupled heterogeneities. We see that the results of the formulae are approximately equal when heterogeneities are relatively small and show that an earlier result in the literature (Koella, 1991) should be viewed in this light. We provide numerical illustrations of our results and discuss a setting in which coupled heterogeneities are likely to have a major impact on the value of *R*_0_. We also describe a rather surprising result: in a system with three heterogeneities, *R*_0_ can exhibit non-monotonic behavior with increasing levels of heterogeneity, in marked contrast to the familiar two heterogeneity setting in which *R*_0_ either increases or decreases with increasing heterogeneity.

## 1. Introduction

The basic reproductive number, *R*_0_, plays a crucial role in determining both whether a pathogen is able to spread and the strength of control measures needed to halt its spread. The simplest descriptions of *R*_0_ assume simple transmission scenarios, such as perfect mixing of a population and homogeneity of the individuals in the population, *e*.*g*. in terms of their susceptibility and infectiousness. The inadequacies of such descriptions have long been realized and much attention has been directed towards understanding the impact of heterogeneities in transmission on the basic reproductive number. Early efforts included accounting for differing activity levels amongst the population and various mixing patterns of the population (*e*.*g*. proportionate/random mixing, assortative and disassortative mixing). Much of this work was prompted by the heterogeneities known to exist for the spread of sexually transmitted infections, notably gonorrhea and HIV (Nold (1980); Hethcote and Yorke (1984); Anderson et al. (1986); May and Anderson (1987); Jacquez et al. (1988); Gupta et al. (1989)). In the context of vector-borne diseases, it has long been realized that vectors’ bites are not distributed uniformly across hosts; instead, there is a heterogeneity in hosts’ attractiveness to vectors, with some individuals being disproportionately favored to receive bites (Carnevale et al. (1978); Dye and Hasibeder (1986); Woolhouse et al. (1997); De Benedicitis et al. (2003); Liebman et al. (2014); Cooper et al. (2019)).

The simplest theoretical framework for exploring the impact of heterogeneities is the multi-type model (Diekmann and Heesterbeek (2000); Diekmann et al. (2012)), in which a population is taken to consist of *n* types (*i*.*e*. epidemiologically relevant categories or groups) of individuals. A now standard argument shows that the basic reproductive number for a multi-type transmission system can be calculated as the dominant eigenvalue of the next generation matrix (Diekmann and Heesterbeek (2000); Diekmann et al. (2012)). The entries *k*_*ij*_ of this *n* by *n* non-negative matrix give the average number of secondary infections of type *i* caused by a type *j* individual in an otherwise entirely susceptible population.

Much attention has been directed towards those special cases of heterogeneous transmission that lead to next generation matrices whose dominant eigenvalue is analytically tractable and hence for which the basic reproductive number can be calculated explicitly. In the context of spatial heterogeneity, these include symmetric spatial configurations such as equally-sized patches with all-to-all or nearest neighbor contacts (see, for example Lloyd and May (1996)).

More generally, a commonly-studied situation involves separable transmission (Diekmann and Heesterbeek (2000); Diekmann et al. (2012)), where the entries of the next generation matrix have the form *k*_*ij*_ = *p*(*i*)*a*(*i*)*b*(*j*) Here, the *a*(*i*) reflect the susceptibility of type *i* individuals, the *b*(*j*) the infectivity of type *j* individuals and the *p*(*i*) give the probabilities that a randomly chosen individual in the population will be of type *i*. The quantities *p*(*i*)*a*(*i*)*/* ∑_*k*_ (*p*(*k*)*a*(*k*)) give the probabilities that an infection caused by a type *j* individual is of type *i*. By definition for separable transmission, these quantities do not depend on *j*. In this case, the next generation matrix is of rank one and has dominant eigenvalue

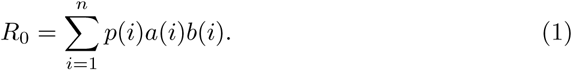

(Note that we chose to write the *p*(*i*) explicitly, whereas most authors’ notation leaves them as a component of the *a*(*i*). One reason for us doing this is that it emphasizes that *R*_0_ does not increase simply as a consequence of there being more groups: increasing the number of groups typically reduces the *p*(*i*) accordingly.)

This expression for *R*_0_ is seen to be an average or expectation taken over the different groups, accounting for the probabilities *p*(*i*), and can be written as

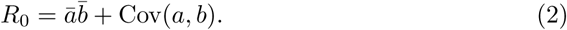

Here, ā and 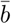 denote the average values of *a*(*i*) and *b*(*i*), and Cov(*a, b*) their covariance. Using the result

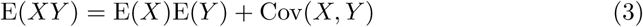

for the expectation of a product of random variables, eqn (1) can be rearranged into the following well-known formula (Dietz, 1980; Dye and Hasibeder, 1986) that sheds insight into the impact of heterogeneity on *R*_0_ in this separable setting:

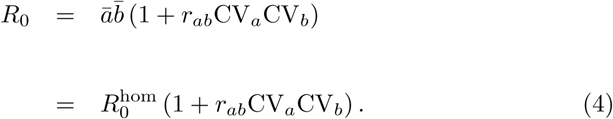

Here, *r*_*ab*_ denotes the Pearson product-moment correlation coefficient between the *a*(*i*) and *b*(*i*),

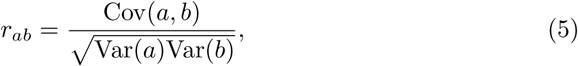

(*i*.*e*. the ratio of their covariance to the product of their standard deviations). CV_*a*_ and CV_*b*_ denote the coefficients of variation (*i*.*e*. standard deviation divided by the mean) of *a*(*i*) and *b*(*i*) and 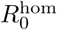 denotes the value of *R*_0_ that would be predicted if the heterogeneity was ignored, *i*.*e*. the average values of *a*(*i*) and *b*(*i*) were used. We emphasize that these results are exact, holding for arbitrary distributions of the *a*(*i*), *b*(*i*) and *p*(*i*).

Heterogeneity can either inflate or deflate the value of *R*_0_, depending on whether there is positive or negative correlation between susceptibility and infectivity across the groups (Dietz, 1980). In the special case where susceptibility and infectivity are proportional, *e*.*g*. for a situation such as differing activity levels or mosquito biting preferences where the heterogeneity impacts both susceptibility and infectiousness in the same way, the formula reduces to

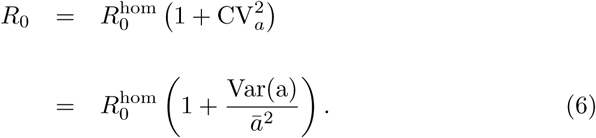

This formula has appeared in the literature numerous times in a number of different settings and guises (Dietz, 1980; Dye and Hasibeder, 1986; May and Anderson, 1987).

Although discussion of the impact of heterogeneity on *R*_0_ is most commonly framed in terms of the multi-type model, similar theory and results can be developed for continuously-distributed heterogeneities (Diekmann et al., 1990; Diekmann and Heesterbeek, 2000; Diekmann et al., 2012). The normal distribution is often used as a first model to describe biological variation and so is a natural description for epidemiological heterogeneity (Sartwell, 1950). However, when variation is moderate or large, *i*.*e*. when the coefficient of variation is not small compared to one, a normal distribution will have substantial weight at negative values. For non-negative quantities, a log-normal distribution can be a more natural choice (Sartwell, 1950; Lessler et al., 2009) that provides a more satisfactory description when variation is not small. Another commonly-used choice is the negative binomial distribution (Lloyd-Smith et al., 2005), with its overdispersion parameter, *k*, allowing it to describe a wide variety of levels of variation from the Poisson through to highly overdispersed. Other descriptions used in models include gamma (Cooper et al., 2019) and power-law distributions. As is usual in mathematical modeling, distributional choices reflect a balance between biological plausibility and mathematical tractability. For the purposes of this study, the continuous formulation is attractive from a mathematical viewpoint because of analytic results that are available for distributions such as the normal and lognormal.

Particularly with the increasing realization that many systems are subject to multiple, often coupled, heterogeneities (Paull et al., 2012; Vazquez-Prokopec et al., 2016), an important question is whether results such as eqns. (4) and (6) generalize to situations in which there are more than two heterogeneities. In this paper, we show that the answer to this question is no: the effect of multiple interacting heterogeneities on the basic reproductive number depends on the details of the distributions of the heterogeneities, in contrast to what occurs in the two-heterogeneity setting. We provide results for both multivariate normal (MVN) and multivariate log-normal (MVLN) distributions of heterogeneities and demonstrate that the two settings can give markedly different results. Furthermore, we show that *R*_0_ can have a non-monotonic relationship with the magnitude of a heterogeneity in a way that does not happen in the two-heterogeneity setting.

## 2. The Model

### 2.1. Discrete Heterogeneities: Multi-type Model

We assume that there are *n* types of individuals, resulting from *N* different heterogeneities, *N*_1_ of which impact susceptibility and *N*_2_ of which impact infectiousness. We write the components of susceptibility as *a*_*x*_ and those of infectivity as *b*_*y*_. An individual’s type can be specified either by their vector of traits, 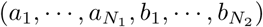, in a *N* = *N*_1_ + *N*_2_-dimensional trait space, or, because we are in an *n*-type group setting, by the index *i* that specifies their group (and hence locates their position within trait space).

We further assume that transmission is separable and that the various heterogeneities impact the susceptibility and infectivity of an individual in a multiplicative fashion. Thus, we assume that the susceptibility of a type *i* individual can be written as the product 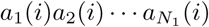, taken over the heterogeneities that impact susceptibility, that the infectivity of a type *j* individual can similarly be written as 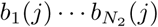, and that the probability that a randomly chosen individual from the population is of type *i* is *p*(*i*).

The entries of the next generation matrix will have the form

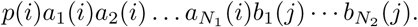

This matrix is of rank one and has dominant eigenvalue given by

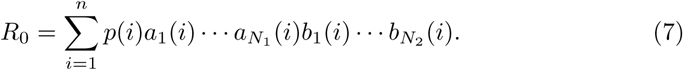

This formula for *R*_0_ can be again be viewed as an expected value, this time of the *N* -way product of the various *a*_*x*_ and *b*_*y*_. Noting that we do not need to distinguish whether a given heterogeneity arises from susceptibility or infectivity when we calculate *R*_0_, we choose to drop the notational distinction between the *a*_*x*_ and the *b*_*y*_, instead referring to a common set *x*_*z*_, where *z* runs from 1 to *N* = *N*_1_ + *N*_2_. This allows us to write common formulae that cover cases where there are a given total number, *N*, of heterogeneities, rather than separate formulae for different combinations of numbers of susceptibilities and infectivities that sum to *N*.

### 2.2. Continuous Heterogeneities

As described above, an analogous model can be formulated for continuously-distributed heterogeneities. A general model for continuous trait space settings is described in (Diekmann et al., 1990; Diekmann and Heesterbeek, 2000; Diekmann et al., 2012). The central component of the model is the kernel function, *k*(*ξ, η*), that gives the average number (per unit volume of trait space) of infections with trait *ξ* that are caused by one individual of trait *η* over their entire period of infectiousness. A map from one generation to the next is given in terms of a next generation operator, which involves integrals over trait space weighted by the kernel. For separable transmission, the kernel can be decomposed as *k*(*ξ, η*) = *p*(*ξ*)*a*(*ξ*)*b*(*η*) and the basic reproductive number can be found as the following integral

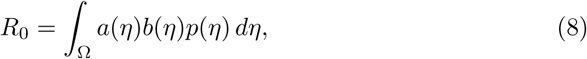

taken over the trait space, Ω. As observed by Diekmann *et al*., while there are some technicalities, in essence—and for our purposes here—moving from the multi-type model to the continuous trait space model involves replacing the discrete sums described in the previous section by integrals.

An individual is specified by their vector of traits 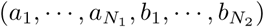 in the *N* = *N*_1_ + *N*_2_-dimensional trait space. In an analogous way to the discrete model, we assume that transmission is separable and that heterogeneities impact susceptibility and infectivity in a multiplicative fashion. This means that the kernel *k* can be written as 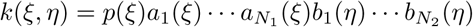 and that *R*_0_ is given by the integral

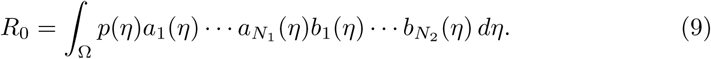

Again, the basic reproductive number can be viewed as the expected value of the product 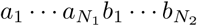 As before we can drop the distinction between the *a*_*x*_ and *b*_*y*_ when performing this calculation: the formula for *R*_0_ reduces to computing the expectation of an *N* -way product of random variables.

## 3. Analysis

The well-known result (eqn 4), as explained above, arises from seeing the formula for the basic reproductive number under separable transmission (eqn 1) as an expectation of the product of a pair of random variables and the ability to express this in terms of the pair’s two expectations and covariance (eqn 3), and hence ultimately in terms of coefficients of variation and the correlation coefficient (eqn 4).

Answering our main question—whether the result can be extended to the case of three or more heterogeneities—requires corresponding manipulations of expectations of products of three or more random variables—the so-called product moments of the joint distribution. We remind ourselves that a crucial part of the standard analysis in the two heterogeneity case was the generality of the formula for the expected value of a product of a pair of random variables, which held for arbitrary distributions.

### 3.1. Product Moments of Random Variables; Main Analytic Results

Closed-form formulae for product moments of multivariate distributions are not generally available. As a result, our analytic results are limited to certain distributions for which such formulae are available. These include the multivariate normal (MVN) and multivariate log-normal (MVLN) distributions, both of which are continuous distributions. To our knowledge, formulae for product moments are not available for widely-used discrete multivariate distributions. Hence, our analytic results focus on continuously-distributed heterogeneities.

In the case of a set of random variables whose joint distribution is multivariate normal, numerous authors have obtained results for product moments (see, for example, Isserlis (1918), Bendat and Piersol (1966), Bär and Dittrich (1971) and Song and Lee (2015)). The general formula for an *N* -dimensional multivariate normal distribution is, however, unwieldy, so we do not present it here. In the four dimensional case, with (*X*_1_, *X*_2_, *X*_3_, *X*_4_) distributed according to a 4-dimensional multivariate normal distribution, we have (Bendat and Piersol (1966) and Bär and Dittrich (1971))

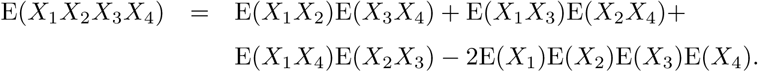

Each expectation of a pairwise product in this equation can be rewritten in the way described above (first using eqn (3) to write in terms of the product of a pair of expectations and their covariance, and then using eqn (5) to rewrite the covariance in terms of coefficients of variation and correlation coefficients) to give

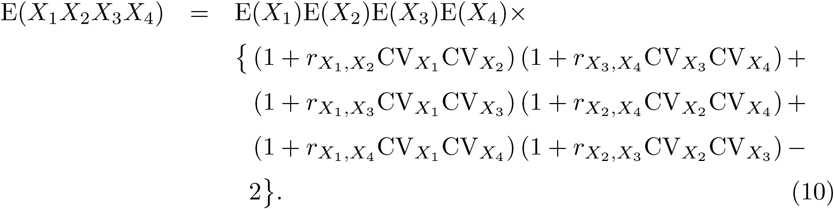

The case of three random variables can be obtained by setting *X*_4_ = 1. In this case, 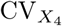 is equal to zero, and so the three parenthetic terms 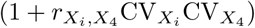 each equal one, leaving

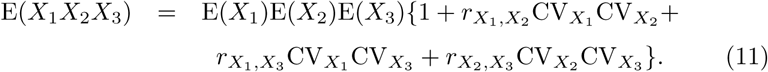

For a set of *N* multivariate lognormally distributed random variables, product moments are given by the formula (Kotz et al., 2000)

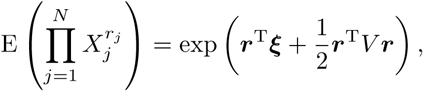

where ***ξ*** and *V* are the mean and variance of the corresponding multivariate normal distribution. In the four dimensional case, some simple manipulation leads to

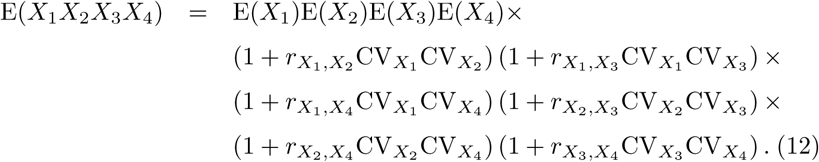

For the case of three random variables, setting *X*_4_ = 1 gives

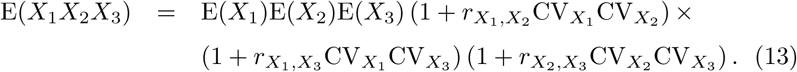

We see that different expressions are obtained for the expectation of a product of four random variables between the multivariate normal (eqn 10) and multivariate lognormal (eqn 12) distributions—for instance, terms in the former involve products of at most two correlation coefficients, whereas the latter includes a term in which all six correlation coefficients are multiplied. Differences are similarly seen for the expectation of a product of three random variables for these two distributions (eqns (11) and (13)).

The observation that eqns (10) and (12) differ, and that their reduced forms when *X*_4_ = 1 also differ, shows that that there is no general formula of this type for the basic reproductive number when there are three or more coupled heterogeneities. This answers the main question posed in this paper. We do notice, however, that the two formulae give approximately equal results in the limit of small coefficients of variation, *i.e*. when one can ignore terms involving products of more than two pairs of coefficients of variation (these are terms that involve products of two or more correlation coefficients). In this case, both eqns (10) and (12) give

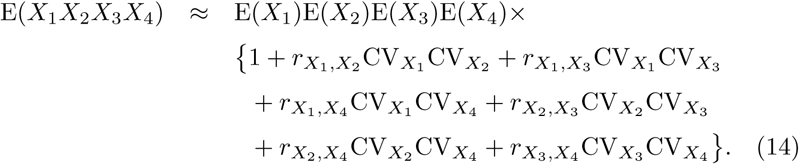

For the case of three random variables, we see that the small coefficient of variation approximation of the three-way MLVN formula (eqn 13) is precisely the formula obtained from the MVN (eqn 11).

### 3.2. Comparison to Previous Results

The majority of papers in the literature that provide analytic results for the basic reproductive number under heterogeneity focus on at most two coupled heterogeneities. One notable exception is the work of Koella (1991), which considers a vector-borne pathogen subject to heterogeneities in mosquito biting rate, *a*, human susceptibility, *b*_2_, and duration of human infection, *ρ*. In a homogeneous setting, the basic reproductive number (strictly speaking, the type reproductive number (Heesterbeek and Roberts, 2007) that considers the complete two-step transmission cycle of the infection) is given by

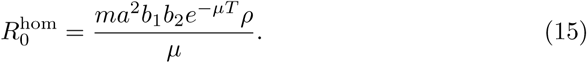

Here, *m* is the ratio of adult female mosquitoes to humans, *b*_1_ is the per-bite transmission probability from human to mosquito, *b*_2_ is the per-bite transmission probability from mosquito to human, *µ* is the mosquito death rate (so that 1*/µ* is the average lifespan of an adult female mosquito) and *T* is the duration of the extrinsic incubation period (*i.e*. the latent period for the infection within the mosquito). Note that the biting rate *a* appears as a squared term, reflecting that the transmission cycle involves two bites by the vector.

Koella provides—without proof or qualification for its applicability—the following formula for the basic reproductive number when *a, b*_2_ and *ρ* exhibit heterogeneity:

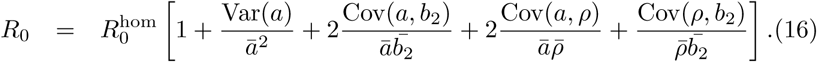

Note that the single biological heterogeneity in biting rate impacts both infectiousness and susceptibility, resulting in it being treated as two perfectly correlated heterogeneities. (The version of eqn (16) given in Koella (1991) appears to have minor but important typographical errors, which are corrected here. In each of the denominators of Koella’s eqn 3, an overbar appears over the entire denominator, whereas overbars should appear separately over individual terms.)

In our notation, we write *X*_1_ = *X*_2_ = *a, X*_3_ = *b*_2_ and 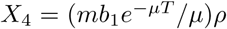. Here *X*_1_ and *X*_3_ impact susceptibility, while *X*_2_ and *X*_4_ impact infectiousness. (We choose to write all of the constant factors within *X*_4_ purely for notational convenience.) We note that eqn (16) has no terms that involve products of pairs of covariances (or, in the language of the earlier formulae, correlation coefficients). Rewriting Koella’s formula in our notation, we see that it follows the approximate formula eqn (14) and so should be seen as an approximation that is likely most accurate when coefficients of variation are small (*i.e*. the heterogeneities are relatively minor). We remark that the Koella formula does agree with the exact result for the multivariate normal distribution, eqn (10), if the coefficient of variation describing either human susceptibility or the duration of human infection is equal to zero (*i.e*. in the case of three coupled heterogeneities).

## 4. Results

### 4.1. Results from Analytic Formulae

We present numerical results obtained from the analytic formulae, allowing us to explore the differences between predictions made using the formulae for the two distributions and also using the formula in the small coefficient of variation limit. For concreteness, we place these simulations within the vector-host setting described above, based on Koella (1991), but for simplicity we hold one of the factors constant. Specifically, mosquitoes differ in their biting rate, which impacts both infectivity and susceptibility and thus is treated as two perfectly correlated heterogeneities, *X*_1_ and *X*_2_, and hosts differ in their duration of infectiousness, *X*_3_. Setting *X*_4_ = 1 and taking *X*_2_ = *X*_1_, we obtain the following two formulae:

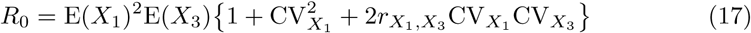

for multivariate normally (MVN) distributed heterogeneities, and

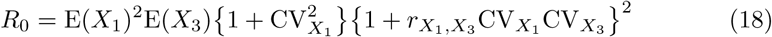

for multivariate lognormally (MVLN) distributed heterogeneities. As discussed above, in this reduced setting of *X*_4_ = 1, the MVN formula is the same as the the small coefficient of variation limit of the MVLN formula and agrees with the Koella formula. Furthermore, we see that when the correlation coefficient, 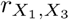, between the two heterogeneities is zero, the two formulae are identical.

Figure (1) compares the *R*_0_ values for heterogeneities distributed according to MVN (solid line, from eqn 17) and MVLN (dashed line, from eqn 18) distributions, across varying levels of correlation between the heterogeneities. We see that the two distributions do indeed lead to identical *R*_0_ values when the correlation coefficient is zero. For positive values of the correlation coefficient, the MVLN distribution leads to a larger value of *R*_0_ than the MVN, while the opposite is true for negative values of the correlation coefficient.

Figure (2) depicts how *R*_0_ values for heterogeneities distributed according to MVN (solid lines) and MVLN (dashed lines) distributions differ across varying levels of correlation between the heterogeneities, for three different levels of heterogeneity in the human duration of infection, *X*_3_. Heterogeneity in *X*_3_ is smallest for the blue lines (Var(*X*_3_) = 1), greatest for the red lines (Var(*X*_3_) = 10), with an intermediate level for the black lines (Var(*X*_3_) = 4: note that these black lines are identical to those in Figure 1). We see that larger levels of heterogeneity lead both to larger deviations in *R*_0_ (increases or decreases, respectively, for positive or negative values of the correlation coefficient 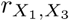) from the value seen when the two heterogeneities are uncorrelated, and larger differences between the values of *R*_0_ that result from use of the two different distributions.

**Figure 1:**
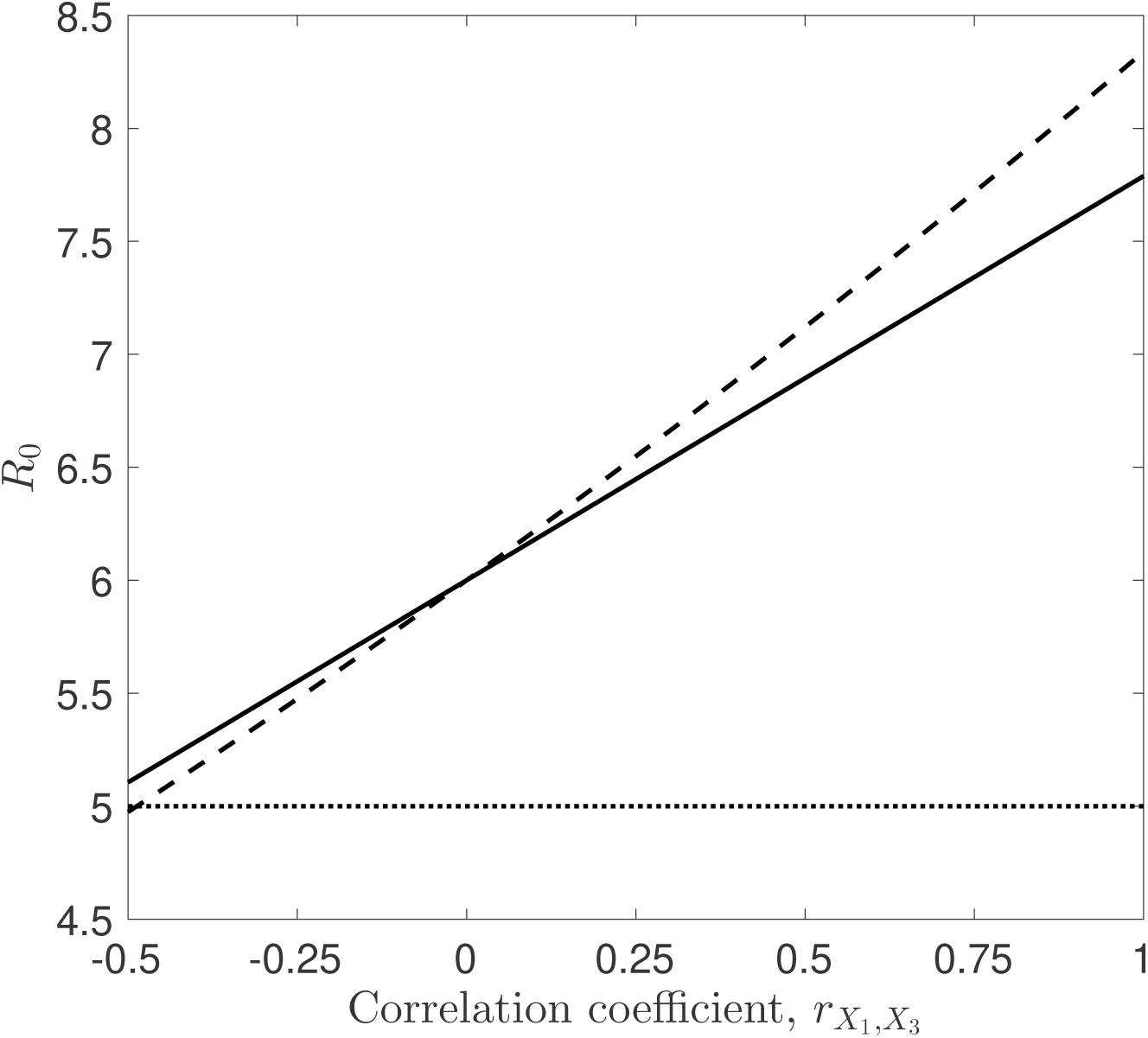
Comparison of *R*_0_ values calculated for multivariate normally-distributed (solid line) and log-normally-distributed (dashed line) heterogeneities for differing values of the correlation coefficient, 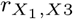, between the biting rate, *X*_1_, and average duration of human infection, *X*_3_. *X*_1_ and *X*_3_ are taken to have means equal to 1 and 5, respectively, and variances Var(*X*_1_) = 0.2 and Var(*X*_3_) = 4. The value of the susceptibility parameter was fixed at 1 for the entire population. Lines are obtained using the MVN and MVLN formulae (eqns 17 and 18, respectively). The horizontal dashed line denotes the value of *R*_0_ if there was no heterogeneity, *i.e*. obtained using the average values, 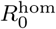. Note that the basic reproductive number is the same for both distributions when the correlation coefficient is equal to zero, and that this is greater than the value 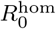 obtained using the average values because of heterogeneity in the biting rate.

**Figure 2:**
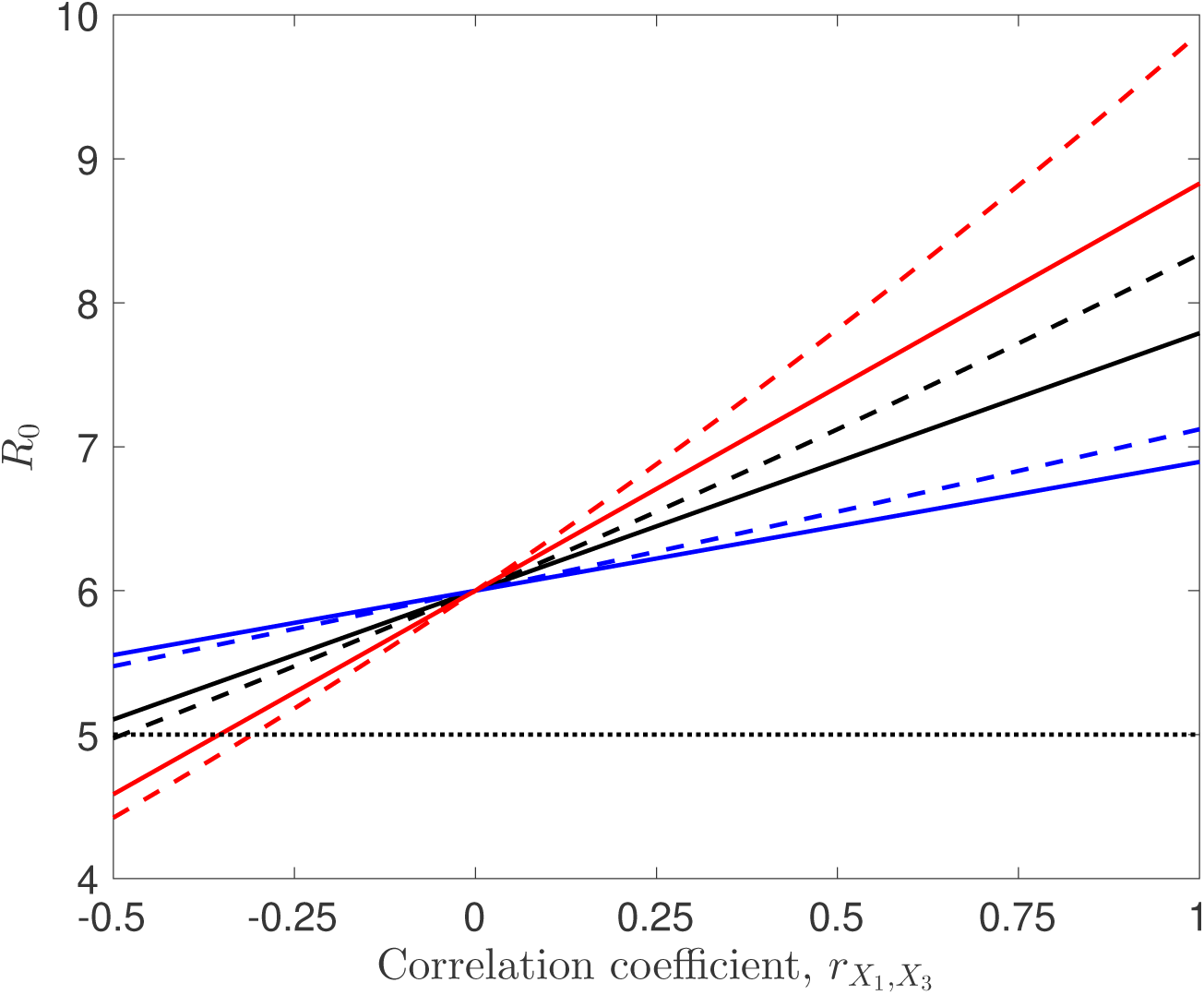
Comparison of *R*_0_ values calculated for multivariate normally-distributed (solid line) and log-normally-distributed (dashed line) heterogeneities for differing values of the correlation coefficient, 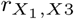, between the biting rate, *X*_1_, and average duration of human infection, *X*_3_. Results are presented for three different levels of the heterogeneity in the average duration of human infection: black lines have Var(*X*_3_) = 4 (these are the lines that appear in Figure (1); blue lines have Var(*X*_3_) = 1, while red lines have Var(*X*_3_) = 10. As in Figure (1), *X*_1_ and *X*_3_ are taken to have means equal to 1 and 5, respectively, and Var(*X*_1_) = 0.2. The value of the susceptibility parameter was fixed at 1 for the entire population. The horizontal dashed line denotes the value of *R*_0_ if there was no heterogeneity, *i.e*. obtained using the average value, 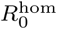.

Figure (3) depicts how *R*_0_ values for heterogeneities distributed according to MVN (solid lines) and MVLN (dashed lines) distributions differ across varying levels of the heterogeneity in biting rate, *X*_1_, for two different levels of correlation between the heterogeneities. Figure (3a) depicts results under a positive correlation, 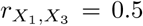, while Figure (3b) shows results for a negative correlation, 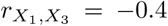. For positive values of the correlation, we see that increasing levels of heterogeneity in biting rate lead to larger values of *R*_0_, with larger values seen under the MVLN than the MVN, in agreement with observations made in earlier figures. Both distributions give the same value of *R*_0_ when there is no variation in biting rate; this equals 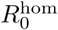 obtained using the average values. Note that *R*_0_ varies nonlinearly with the level of variation in biting rate, reflecting the biting rate’s quadratic influence on *R*_0_.

**Figure 3:**
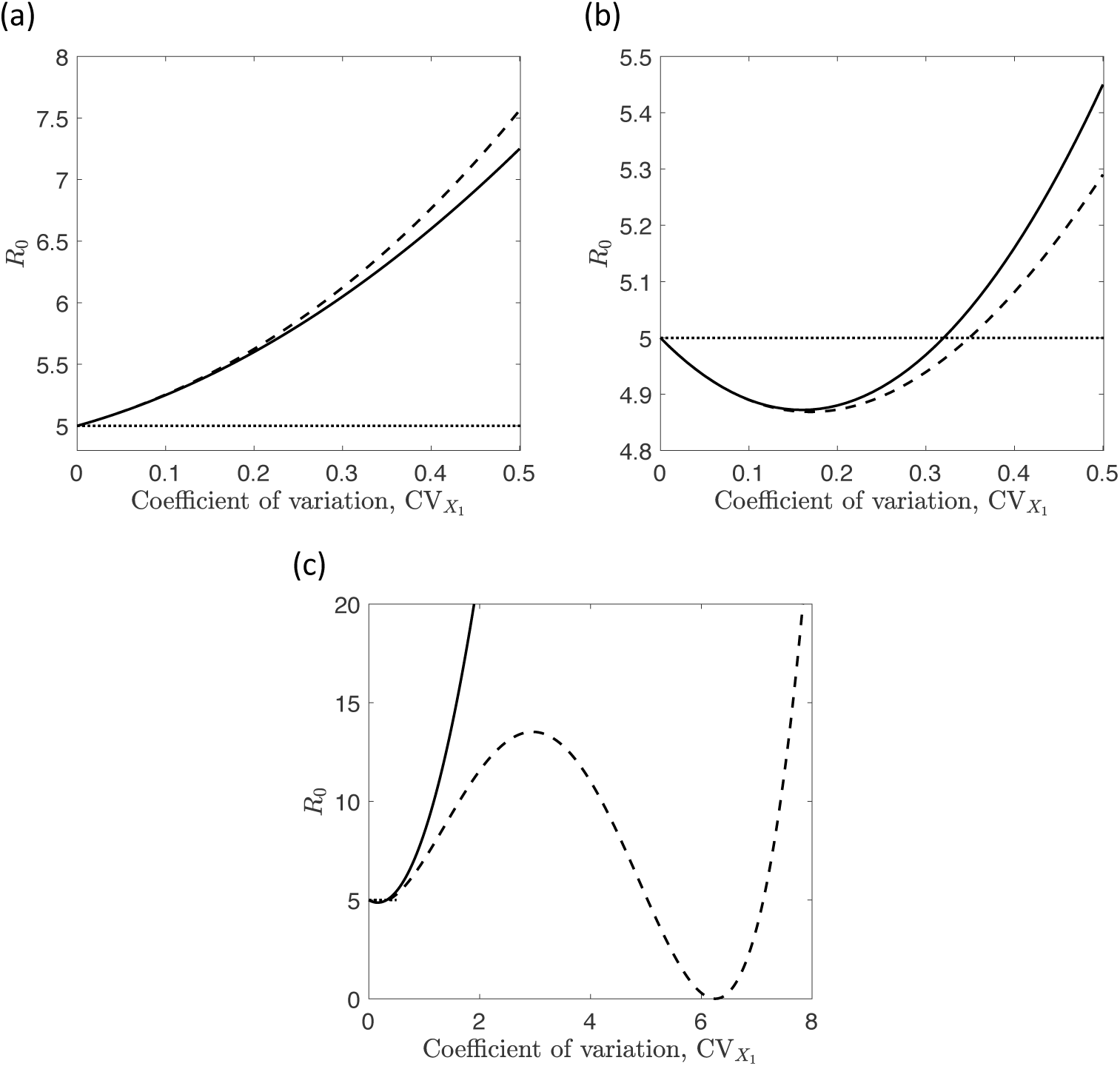
Comparison of *R*_0_ values calculated for multivariate normally-distributed (solid line) and log-normally-distributed (dashed line) heterogeneities for differing levels of heterogeneity in the biting rate, *X*_1_. Panel (a): positive correlation coefficient, 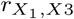, between the biting rate and average duration of human infection, *X*_3_, equal to 0.5. Panel (b): negative correlation coefficient, with 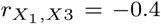. Panel (c): as panel (b), but with a larger range of values of CV(*X*_1_) on the horizontal axis. Other parameter values are as in Figure 1, with *X*_1_ and *X*_3_ taken to have means equal to 1 and 5, respectively, and Var(*X*_3_) = 4. The value of the susceptibility parameter was fixed at 1 for the entire population. The horizontal dashed lines denote the value of *R*_0_ if there was no heterogeneity, *i.e*. obtained using the average value, 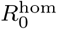. Note the surprising non-monotonic impact of heterogeneity in panels (b) and (c), behavior that does not occur in the well-studied case of two coupled heterogeneities.

Figure (3b) provides a very surprising result: when the correlation coefficient is negative, the basic reproductive number can exhibit non-monotonicity as the level of heterogeneity increases. This result, which does not occur in the two heterogeneity setting, arises from *R*_0_ increasing in a quadratic manner on the heterogeneous biting rate but decreasing in a linear fashion on the interaction between heterogeneities in biting rate and duration of infection. Nonmonotonicity will always occur as the magnitude of a heterogeneity is varied if that heterogeneity impacts both susceptibility and infectivity and is also negatively correlated with another heterogeneity.

Figure (3c) shows that for the MVLN distribution, non-monotonicity can be more severe than depicted in Figure (3b). For this distribution, *R*_0_ depends in a quartic manner on the magnitude, 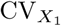, of the heterogeneity *X*_1_, and so there can be multiple local minima. Looking over the larger range of values for 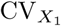 shown in this panel (except for this, all other details are as in panel b), one sees a local maximum in *R*_0_ when 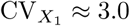 and a second local minimum when 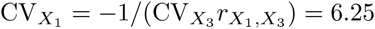. This second local minimum, at which *R*_0_ is equal to zero, arises when the term in braces that gets squared in eqn (18) equals zero.

In the four heterogeneity formulae, setting *X*_1_ = *X*_2_ gives *R*_0_ as a quadratic function of 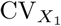 in the MVN case and a sixth degree polynomial in the MVLN case.

### 4.2. Multi-Type Model Results

We conclude with a numerical example in a multi-type setting, based on real-world data from Cooper et al. (2019) on heterogeneity in mosquito biting rates across groups in a study in Uganda. Cooper *et al*. fit gamma distributions to the relative numbers of bites received by different groups (for them, households) for a number of locations. The degree of heterogeneity was found to vary across location and time; we use the gamma distribution fitted to relative bite count data from Jinja District, for which the standard deviation was 0.985 and, by definition, the mean was 1. Note that the coefficient of variation here is large compared to values depicted in Figures (3a and b).

We employ a thousand group model, with an equal number of individuals (people) in each. We assume that groups differ in their attractiveness to mosquitoes and in the average duration of human infection. By definition in this multi-group model, all members of a group are taken to have identical attributes. We write the relative attractiveness and duration of human infection of group *i* as *z*_*i*_ and *ρ*_*i*_, respectively. As in the Cooper *et al*. study, we take the *z*_*i*_ to have mean one, so that the per-mosquito biting rate on group *i, a*_*i*_, is given by the product of the average biting rate and the relative attractiveness: *a*_*i*_ = *āz*_*i*_. Finally, we assume some level of correlation, *r*_*a,ρ*_, between the *z*_*i*_ (or, equivalently, the *a*_*i*_) and *ρ*_*i*_. Because of the lack of data on multiple coupled heterogeneities, we will explore the impact of different levels and signs of correlation between the two variables.

We base our parameter values on those used by Chitnis et al. (2008) in a modeling study of malaria transmission, using their high intensity transmission scenario. We take the average mosquito biting rate, *ā*, to be 0.5 per day, the ratio of vectors to hosts, *m*, to be 9.27, the per-bite transmission probabilities *b*_1_ and *b*_2_ to equal 0.022 and 0.48, respectively, the average mosquito lifespan, 1*/µ*, to be 10 days, and the extrinsic incubation period, *T*, to be 11 days. We assume the duration of human infection is lognormally distributed, with mean 290 days (corresponding to the 9.5 month average duration described by Chitnis *et al*.) and coefficient of variation 0.4.

For each group, we draw the relative biting attractiveness, *z*_*i*_, and duration of human infection, *ρ*_*i*_, from a bivariate distribution with gamma and lognormal marginals as described above. These numbers are obtained using the NORTA algorithm (Cario and Nelson, 1997, 1998), as implemented by the NORTARA package (Su, 2014) in R (R Core Team, 2019). This algorithm allows generation of random numbers from multivariate distributions with specified marginal distributions and correlation structure.

The next generation matrix (considering the complete transmission cycle from human to human, via mosquito) has entries *k*_*i j*_, where

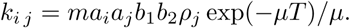

We see that this next generation matrix is separable and involves the product of three group-dependent quantities (*a*_*i*_, *a*_*j*_ and *ρ*_*j*_).

If there was no heterogeneity, the value of the basic reproductive number would equal 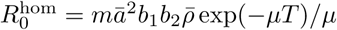, and take the value 23.63 given the parameters employed here. If the only heterogeneity was in mosquito biting preference, the basic reproductive number would be inflated by the factor 1 + Var(*z*_*i*_) = 1.97, giving an *R*_0_ of 46.58. (The attentive reader might notice that our values of *R*_0_ appear quite different to those of Chitnis et al. (2008) even in the homogeneous setting. The major difference arises because we calculate values on a per transmission cycle basis rather than the per transmission event basis employed by Chitnis et al. (2008): their *R*_0_ formula is essentially the square root of the one used here. Beyond this, there are some minor differences in model formulations, such as our use of a fixed duration of latency.)

For a given set of values of *a*_*i*_ and *ρ*_*j*_, the value of *R*_0_ for this model with its coupled heterogeneities can be calculated exactly using eqn (7). We would like to be able to relate this to the moments of the individual heterogeneities, but we do not have a closed form expression for the product moments of the multivariate distribution employed in this example. As we are in a three heterogeneity setting, we could use the formulae derived using the multivariate normal (eqn 17) or multivariate lognormal (eqn 18) distribution. Both will be approximations because they were derived using distributions different to the one employed in this example. Recall that in this three heterogeneity setting, the MVN formula is identical to the small coefficient of variation limit of the MVLN formula and to the Koella formula. (It should be kept in mind, however, that coefficients of variation are large in this numerical example, meaning that we are not in the weak heterogeneity limit.) Finally, when using the MVN and MVLN formulae, we can either use means and coefficients of variation calculated based on the set of values of *a*_*i*_ and *ρ*_*j*_, or those of the underlying distribution from which these values were drawn. Because we are using a sampled set of values, moments calculated from the sample will generally differ from those of the underlying distribution.

We remark that the relevant product moment of the continuous multivariate gamma/lognormal distribution that underlies this example (i.e. the distribution from which the groups’ attributes are sampled) could be obtained numerically using Monte Carlo numerical integration. Indeed, we see that the value of *R*_0_ obtained from eqn (7) can be viewed in just this way as an (admittedly crude) Monte Carlo estimate. One approach to obtain a better approximation would be to repeat the group attribute sampling process, evaluating *R*_0_ each time, and averaging the resulting collection of estimates. Alternatively, one could obtain a more representative sample of the underlying distribution by simply increasing the number of groups.

Results are shown in Table (1), and are as anticipated from the discussion of the analytic results. As mentioned above, heterogeneity in biting alone inflates the value of *R*_0_ to 46.58, above the value of 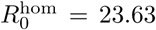 that would be predicted if each group’s attributes were simply equal to the average of the underlying distribution. Exact results obtained from the 1000 group model (the column labeled “Numerical (Exact)”), show that positive correlation between the biting attractiveness and duration of infectiousness further increases *R*_0_, with a greater increase when the correlation is stronger. Negative correlation decreases the value of *R*_0_. Note that the exact value obtained when the correlation is zero differs from the 46.58 stated above: sampling variation (when drawing the 1000 sets of attributes) means that the averages of the *a*_*i*_ and *ρ*_*j*_ for the 1000 groups differ from those of the underlying distribution.

**Table 1:** *R*_0_ values for the multi-group host-vector transmission model. As described in the text, the groups differ in terms of the mosquito biting rate (*a*) and the duration of human infection (*ρ*), with different levels and directions of correlation between the two heterogeneities (*r*_*aρ*_). The “Numerical (exact)” column gives (exact) numerical results obtained from a 1000 group model, with group attributes sampled from a bivariate distribution with relative biting attractivenesses of the groups given by a gamma distribution with mean 1 and standard deviation 0.985 (Cooper et al., 2019) and durations of human infection lognormally distributed with mean 290 days and standard deviation 116 days. Other parameters are given in the text and lead to a value of 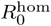 of 23.63. MVN and MVLN formulae refer to results obtained using analytical formulae (eqns 17 and 18), either using means and coefficients of variation obtained from the set of attributes in the 1000 group model, or by using the corresponding quantities from the underlying continuous distribution from which group attributes are sampled. Monte Carlo refers to results obtained using numerically estimated product moments of the underlying continuous distribution from which group attributes are sampled. Monte Carlo estimates were obtained by averaging over 4000 sets of results for the 1000 group model. Both the standard error (quoted as *±* value) of this estimate and the standard deviation across the 4000 sets of results (value in parentheses) are given. The standard error, which describes the uncertainty in the Monte Carlo estimate, is obtained by dividing the standard deviation by 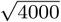. The standard deviation describes the dispersion of *R*_0_ values obtained from single realizations of the 1000 group model about the underlying mean.

| $r_{a,\rho}$ | Sample | | | Underlying Distribution | | |
| --- | --- | --- | --- | --- | --- | --- |
|  | Numerical<br>(Exact) | MVN<br>formula | MVLN<br>formula | Monte Carlo | MVN<br>formula | MVLN<br>formula |
| 0.8 | 75.89 | 65.56 | 84.17 | $73.58 \pm 0.12$ (7.54) | 61.48 | 80.58 |
| 0.5 | 54.62 | 50.66 | 58.54 | $62.90 \pm 0.09$ (5.81) | 55.89 | 66.74 |
| 0.0 | 49.15 | 48.47 | 49.14 | $46.60 \pm 0.06$ (3.56) | 46.58 | 46.58 |
| -0.5 | 38.15 | 45.27 | 35.49 | $32.49 \pm 0.03$ (1.88) | 37.26 | 30.03 |

Values obtained from the single realization of the 1000 group model are consistent with the Monte Carlo estimates of the basic reproductive number that would pertain if the heterogeneity exactly followed the underlying continuous multivariate distribution. Some variation from these values is observed: this is a consequence of the attributes in the 1000 group model being a not so large sample drawn from a distribution with marked variation and skewness. We see that the MVLN formula provides a closer approximation to the exact value of the reproductive number than does the MVN formula (which we recall is identical to both the small heterogeneity and Koella’s formula). The MVLN formula does tend to overestimate the impact of the coupled heterogeneity, in each case moving it further away (either in a positive or negative direction, depending on whether *r*_*a,ρ*_ is positive or negative) from the value obtained with zero correlation, compared to the Monte Carlo estimate.

To emphasize the difference between settings involving three or more heterogeneities and the more familiar two heterogeneity setting, we remind the reader that equation (4) for *R*_0_ is exact in the two heterogeneity case, regardless of the details of the distributions of the heterogeneities. Values in the three “Sample” columns of a corresponding table in the two heterogeneity case would be equal, as would be the values in the two MVN and MVLN “Underlying Distribution” columns.

## 5. Discussion

In this paper we have shown that the well-known result for the impact of two coupled heterogeneities on the basic reproductive number of an epidemiological system under separable transmission does not have a general counterpart when there are three or more coupled heterogeneities. In the more general setting, the formula for the basic reproductive number depends on details of the joint distribution of the heterogeneities in a way that is quite different than in the setting with two heterogeneities. We were able to derive formulae that related the basic reproductive number to the magnitudes of the heterogeneities and their pairwise correlations for the special cases of multivariate normal and multivariate lognormal distributions of heterogeneities. Under particular limiting cases (typically in the limit of low levels of heterogeneity), the two formulae give similar predictions. We showed that an earlier result in the literature (Koella, 1991) should be viewed as an approximate result, although we noted that in appropriate limiting cases, the result agrees with our formula for the multivariate normal distribution.

The *R*_0_ formulae obtained here were derived for continuous multivariate distributions and thus can only provide an approximation for discrete multigroup settings such as our numerical example. (Furthermore, our numerical example used an underlying distribution that differed from the ones for which we derived our formulae.) This approximation would improve as the number of groups increased and the multi-group model more closely approximated a setting in which heterogeneities could be treated as being continuously distributed. This again is in contrast to the two heterogeneity setting, in which the well-known result is exact, applyng to all distributions.

It is only possible to obtain analytic formulae for the basic reproductive number in multi-type models whose structure permits evaluation of the dominant eigenvalue of the next generation matrix. Although separable transmission provides one class of models for which this is the case, and has thus received much attention in the theoretical literature, it should be realized that most real-world transmission settings that involve structure do not fall in this class. Spatial models will almost invariably lead to non-separable transmission: interactions are more likely with nearby than distant individuals, so numbers of encounters depend on the relative locations of both individuals, rather than separately on their two locations. Another example is provided by age structure, commonly discussed in the context of childhood infections. Here, individuals are more likely to interact with individuals of a similar age than those of a different age. In models for the spread of sexually transmitted infections, it is often the case that individuals are more likely to interact with ones who are similar to themselves (*e.g*. highly sexually active people are more likely to have such a partner). Each of these departures results from some form of assortative mixing, a commonly-observed property in real-world social networks Newman (2003). While separable transmission might not be reflective of all real-world settings, that it allows us the ability to gain analytic insights into the impact of heterogeneity means that it remains a useful situation to study, although the limitations of its applicability must be borne in mind.

Although theoretical attention has typically focused on the two heterogeneity case, and this has provided much insight, heterogeneous transmission in the real world typically involves more than two factors (Paull et al., 2012; Vazquez-Prokopec et al., 2016). Vector borne diseases provide striking examples of systems impacted by multiple, coupled heterogeneities (Koella, 1991; Smith et al., 2014; Vazquez-Prokopec et al., 2016; Irvine et al., 2018). Both host and vector populations themselves exhibit epidemiological heterogeneities, such as host movement patterns (Stoddard et al., 2013), duration and severity of host infectiousness (Nguyet et al., 2013) and vector density (LaCon et al., 2014), as well as having significant heterogeneity in their interactions, for instance due to host biting preference and spatial co-location between hosts and vectors (Liebman et al., 2014; Cooper et al., 2019). As such, epidemiological systems such as the human/*Aedes aegypti* /dengue virus or human/*Anopheles*/malaria systems represent prime examples for the application of results such as those discussed here. Our ability to apply these results at present is, however, hindered by the lack of datasets in which multiple heterogeneities have been simultaneously characterized.

When transmission is heterogeneous, control measures that account for the heterogeneity are much more effective than ones that do not (Dye and Hasibeder, 1986; Anderson and May, 1991; Woolhouse et al., 1997; Lloyd-Smith et al., 2005; Cooper et al., 2019), an observation that has driven much of the research into heterogeneity in transmission. The most striking result of this study is the observation that *R*_0_ can have a non-monotonic dependence on the magnitude of a heterogeneity when there are three or more coupled heterogeneities—in marked contrast to the behavior seen in the two heterogeneity case. Indeed, in the case of a multivariate lognormal distribution, there is the potential for multiple local minima and maxima in the basic reproductive to occur as the level of heterogeneity is changed. Non-monotonicity could have important implications for the impact of targeted controls, which typically lead to a reduction in the degree of heterogeneity across the population. In such cases, non-monotonicity could conceivably and perversely lead to an increase in the reproductive number for some levels of control. We comment that the example in which we demonstrated non-monotonicity involved varying the level of heterogeneity while keeping the mean transmission potential constant, while control would involve decreasing both mean and variation. As such, the potential for this phenomenon to cause perverse outcomes of targeted control is not immediate; we defer exploration of this to a later study.

The results of this paper demonstrate the importance of gaining understanding of how multiple coupled heterogeneities impact transmission. Even with the addition of just one additional heterogeneity, our analysis reveals that surprising behavior can be observed beyond what has previously been seen in the setting of two heterogeneities. While providing a theoretical step in this direction, this study highlights the limitations of general results that can be obtained in more realistic settings. It aims to guide more detailed studies that involve numerical exploration of specific situations, yielding further insights into the epidemio-logical role of individual variability. We argue for the need to move beyond quantification of single heterogeneities, to simultaneous measurement of multiple heterogeneities and their covariation, in order to better understand the impact of individual variability on transmission dynamics of infectious diseases.

## Acknowledgments

This work was supported by grants from the National Institutes of Health (P01-AI098670; all authors, and R01-AI091980; ALL) and the National Science Foundation (RTG/DMS—1246991; ALL). Support from the Drexel endowment, NC State University, is also acknowledged by ALL. The authors declare that they have no conflicts of interest. We thank the referees for their helpful comments and probing questions that pushed us to sharpen the presentation of this work.

